# My Gut Feels Your Pain - The Social Transfer of Pain Remodels the Gut Microbiome

**DOI:** 10.64898/2026.01.13.699329

**Authors:** Marisol I. Dothard, Parnaz Boroon, Sabrena C. Tuy, Jie Zhang, Sarah Allard, Monique L. Smith, Jack A. Gilbert

**Affiliations:** Department of Pediatrics and Scripps Institution of Oceanography, University of California San Diego, La Jolla, California, 92093, USA; Biomedical Sciences Graduate Program, University of California San Diego, La Jolla, California, 92093, USA; Department of Neurobiology, University of California San Diego, La Jolla, California, 92093, USA; Department of Neurosciences, University of California San Diego, La Jolla, California, 92093, USA; T. Denny Sanford Institute for Empathy and Compassion, University of California San Diego, La Jolla, California, 92093, USA

## Abstract

**Background:** The “social transfer of pain” is a phenomenon where a mouse experiencing injury-induced hyperalgesia can trigger hyperalgesia in a mouse briefly housed in the same environment (‘bystander’). The peripheral mechanisms underlying social transfer of pain in mice are not yet well described. As gut microbes are associated with social interactions, pain states, and pain attenuation via the gut-brain axis, we hypothesized that the bystander gut microbiome may respond to the social transfer of pain.

**Methods:** To induce the social transfer of pain, complete Freund’s adjuvant (CFA) was injected into the hindpaw of a mouse which then underwent social interaction with a bystander for one hour. Mechanical sensitivity was assessed using the Von Frey mechanical sensitivity test. Stool samples and mechanical thresholds were taken prior to social interaction, 4 hours post-social interaction, and 24 hours post-social interaction. Metagenomic sequencing characterized the taxonomic and predicted functional gene response of the gut microbiome to CFA-induced pain, social transfer of pain and control groups.

**Results:** At 4 hours post-social interaction, bystander animals experienced increased mechanical sensitivity comparable to CFA-injected animals and significantly lower than controls, indicating enhanced hyperalgesia. Compared to baseline, fecal community composition analyses at 4 hours and 24 hours post-interaction showed significant differences in Unweighted Unifrac in both CFA-injected and bystander animals but not in controls. Differential abundance analyses using Maaslin2 identified significant increases in the relative abundances of short chain fatty acid producing taxa like *Lachnospiraceae*, *Ruminococcaceae*, *Oscillospiraceae* and decreases in commensal mouse gut microbes like *Muribaculaceae* from baseline to 4 hours and 24 hours post-interaction in both bystanders and CFA-injected animals but not in controls. Functional analysis revealed increased abundance of pathways related to short-chain fatty acid production including pyruvate to butanoate and L-lysine fermentation to acetate and butanoate. The altered gut microbiome of bystanders strongly resembles that observed in CFA-injected animals at 4 hours and 24 hours post-injection, with the addition of a unique bystander bloom in several species of *Lachnospiraceae*.

**Conclusions:** The changes in the gut microbiome of bystander animals suggest that the social transfer of pain alters bystander peripheral physiology. These results are the first evidence of the potential for such a link.

## Introduction

A growing body of work across animal models and humans demonstrates that the gut microbiome both shapes and is shaped by social behavior, influencing neurodevelopment, stress responsivity, affect, and social cognition. Experimental studies in rodents have shown that manipulation of the gut microbiota alters social interaction, anxiety-like behavior, and stress-induced neuroinflammatory responses, implicating microbial metabolites and immune signaling as mediators of gut–brain communication (Cryan and Dinan 2012; Sherwin et al. 2016). In parallel, comparative and ecological studies in primates and humans reveal that social relationships and network structure strongly predict microbiome composition (Dasari et al. 2025), leading to the concept of a “social microbiome” in which microbial transmission occurs along social ties and may contribute to socially patterned health outcomes (Sarkar et al. 2020, 2024). In humans, microbiome-associated inflammatory markers and specific taxa have been linked to variation in social cognition, emotional processing, and vulnerability to depression, suggesting that microbiome–immune–brain interactions may influence empathy-relevant psychological traits (Heym et al. 2019). Together, these findings raise the possibility that socially transmitted physiological states—including pain—may engage not only neural circuits but also systemic pathways involving immune signaling and the gut microbiome.

Recent studies report a phenomenon in which rodents exhibit pain, analgesia, or inflammation matching the state of a social partner following short-term exposure (Lannon et al. 2024; Castany et al. 2024; Smith et al. 2016, 2021). In one widely used procedure, the “social transfer of pain” can be induced in a social partner (‘bystander’) following a one-hour social interaction with a social partner injected with complete Freud’s adjuvant (CFA). After this social interaction, the bystander demonstrates increased mechanical and thermal sensitivity (Smith et al. 2021; Rein et al. 2022). Mechanical sensitivity testing indicates that bystander mice experience hypersensitivity for more than 4hrs following interaction, with recovery being between 4-24hrs (Smith et al. 2021). Mechanistically, anterior cingulate cortex pyramidal neurons form direct, monosynaptic projections to the nucleus accumbens core, and activation of this pathway is required for the acquisition of socially transferred pain: inhibiting ACC to NAc terminals during a brief social interaction with a mouse in pain prevents the development of hyperalgesia in the bystander animal, whereas optogenetic activation of this pathway prolongs the duration of the transferred pain state.(Smith et al. 2021) However, the response to the pain of a social partner is likely systemic, particularly across the gut-brain axis, which connects the nervous system to the immune system and the gut microbiome. It is unknown whether the gut microbiome changes in response to the social transfer of pain, but as gut microbial metabolism has been shown to play a role in host physiological responses, it is possible that the microbiome may influence the host’s pain phenotype (and vice versa).

To date, only one study has examined links between the gut microbiome, inflammation, and psychological traits relevant to empathy and depression risk in humans, which demonstrated that *Lactobacillus* abundance was positively associated with positive self-judgement and indirectly related to cognitive empathy, while C-Reactive Protein levels are negatively associated with cognitive empathy; however, no pain-sharing or social transfer of pain was measured. (Heym et al. 2019) Given that empathy is a core social process supporting the transmission of distress between individuals, these associative human data highlight the potential for microbiome–brain interactions to influence empathy-related behaviors such as the social transfer of pain. However, because the study examined depressive risk and self-reported empathy rather than behavioral empathic responses or sensory changes, preclinical controlled experiments remain essential to establish whether and how the microbiome modulates neural and behavioral mechanisms of social pain transmission.

Several lines of evidence suggest that the gut microbiome may respond to or even play a role in the mechanisms of the social transfer of pain. For example, evidence suggests that the gut microbiome contributes to pain pathologies through both indirect inflammatory mechanisms and direct activation of sensory neurons. Dysbiosis has been linked to chronic pain states via immune modulation and altered gut–brain signaling (Ustianowska et al. 2022), while recent studies show that bacterial proteases can directly activate nociceptors by engaging protease-activated receptors, providing a microbial route to pain generation (Altier 2024).Further, when CFA is used to induce chronic pain in mice, the gut microbiome demonstrates a reduction of anti-inflammatory short chain fatty acid (SCFA) producing microbes as well as a reduction in SCFAs 4 days after injection. (Ma et al. 2020) The gut microbiome is both altered by and can causally mediate changes in affect, as evidenced by the bidirectional relationship with affective disorders like depression (Naseribafrouei et al. 2014; Zheng et al. 2016; Fan et al. 2022) We therefore hypothesized that the gut microbiome may respond to the social transfer of pain. Given that the social transfer of pain results in matching behaviors between social partners, we further hypothesize that the gut microbial composition of the bystander animals will strongly resemble that found in the gut microbiome of CFA-injected animals.

The aim of this study was to characterize the response of the bystander gut microbiome to the social transfer of pain. We performed taxonomic and functional metagenomic analyses on fecal samples of bystander animals, CFA-injected animals, and control animals prior to, and 4 hours, and 24 hours post social interaction between bystander and CFA-injected animals.

## Methods

### Animals

A total of 48 male adult C57BL/6J mice (8 weeks of age) were obtained from the Jackson Laboratory (Bar Harbor, ME, USA). Mice were delivered at 7 to 8 weeks of age, housed 4 per cage, and acclimated to our colony room under a reversed light/dark cycle (lights off at 07:00, on at 19:00). Behavioral testing was conducted in dedicated behavior rooms under red illumination during the dark phase. These mice were housed in disposable (Innorack IVC Mouse 3.5; Innovive) in a temperature-and humidity-controlled environment with ad libitum access to food (Irradiated Teklad Diet; Inotiv). All procedures were approved by the Institutional Animal Care and Use Committee at University of California San Diego (Protocol No. S22191) and were conducted in accordance with the National Institutes for Health Guidelines for the Care and Use of Laboratory Animals and the Guidelines for the Care and Use of Mammals in Neuroscience and Behavioral Research.

### Behavioral Assays and Stool Collection Methods

Full protocols describing the social transfer of pain paradigm are detailed in Rein et al. 2022. Briefly, 48 C57BL/6J mice were habituated to the testing room and mechanical testing rack the 2 days prior to the social transfer. On the final day of acclimation, baseline mechanical thresholds were acquired and used to counterbalance mice into one of three treatment groups with matched basal mechanical thresholds: control, CFA-injected, or bystander. Baseline stool samples were also taken on the final day of acclimation. Each animal was also assigned a partner within its cage, thereby forming pairs. Each pair included a demonstrator mouse which would exhibit pain via hindpaw injection of 15uL Complete Freund’s Adjuvant (CFA, Sigma F5881) and a bystander mouse (which would interact with the CFA-injected mouse). Control mice were paired together and did not experience any pain induction (control + control). Thus, within each cage of 4, mice were assigned as either 2 pairs of control mice, or 2 CFA-injected and 2 bystander mice. Mechanical thresholds were assessed using the Up-Down technique (Chaplan et al.,1994) prior to any treatment and baseline stool collections were conducted simultaneously. 24 hours later, mice were habituated to the testing room, and CFA-injected mice were injected immediately before the social interaction. CFA-injected mice were then paired with their fellow bystander mouse and placed together in a clean, unfamiliar cage for a one-hour social interaction. Immediately following the social interaction, mechanical sensitivity testing was performed. Pairs were separated and housed within treatment-matched and familiar cages following social interaction. Mechanical sensitivity measurements as well as stool collections were taken at 4 hours and 24 hours post social transfer of pain.

### Metagenomic Sequencing and Analyses

#### Sequencing

The University of California Microbiome Core performed nucleic acid extractions using previously published protocols (Marotz et al. 2021) and 143 fecal samples were processed from bystander animals, CFA-injected animals, and negative control animals. Briefly, extractions and purifications were performed using the MagMAX Microbiome Ultra Nucleic Acid Isolation Kit (Thermo Fisher Scientific, USA) and automated on KingFisher Flex robots (Thermo Fisher Scientific, USA). Blank controls and mock communities (Zymo Research Corporation, USA) were included per extraction plate, which were carried through all downstream processing steps. Input DNA was quantified using a PicoGreen fluorescence assay (Thermo Fisher Scientific, USA) and metagenomic libraries were prepared with KAPA HyperPlus kits (Roche Diagnostics, USA) following manufacturer’s instructions and automated onEpMotion automated liquid handlers (Eppendorf, Germany). Sequencing was performed on the Illumina NovaSeq 6000 sequencing platform with paired-end 150 bp cycles at the Institute for Genomic Medicine (IGM), UC San Diego.

#### Quality filtering and data preparation

Raw metagenomic sequences were quality filtered and trimmed using fastp v 0.22 (Chen et al. 2018), and mouse and PhiX reads were filtered using bowtie2 version 2.22 (Li 2018) and converted to FASTQ format using SAMtools version 1.15.1(Li et al. 2009). Reads were aligned against the Web of Life database (Zhu et al. 2019) using Bowtie2 (Langmead and Salzberg 2012) and classified with the Woltka pipeline (Zhu et al. 2019), resulting in a table of Operational Genomic Units (OGUs). Samples were filtered using a 1% coverage filter.

#### Metagenomic QC and Diversity Analyses

Sequencing controls were removed after performing quality checks. Nine samples (3 animals) were triaged after rarefaction to a common sequencing depth was set at 330,000 sequences per sample, based on rarefaction curves. Shannon Diversity (Russell 1980) was compared within groups longitudinally using linear mixed effect modeling (Shannon ∼ time_point + cohort + (1|mouse_id)). Differences in bacterial community composition were calculated between sample groups using Weighted and Unweighted UniFrac distance matrices (Lozupone Catherine and Knight Rob 2005), and statistical differences were assessed with pairwise or non-pairwise permutational multivariate analysis of variance (PERMANOVA). Differential abundance was performed using the package Maaslin2, with significant results as q < 0.05. version 2.4.0 (Lin and Peddada 2020). Figures were generated in R version 4.3.2 (R core team, 2022) with phyloseq (McMurdie and Holmes 2013), tidyverse (Wickham et al. 2019) and ggplot2 (Wickham 2011) packages.

#### Functional Analysis

Functional profiling was performed using Woltka v0.1.4 with paired-end metagenomic reads aligned against the Web of Life reference protein database (WoLr2) via Bowtie2 v2.4.2 using --very-sensitive mode and permissive scoring (-k 16 --np 1 --mp 1,1 --rdg 0,1 --rfg 0,1--score-min L,0,-0.05). Reads were mapped to protein sequences and collapsed to MetaCyc pathways using the --map and --names flags. Following classification, pathway abundance tables were filtered using a custom Python script based on prevalence and abundance thresholds to remove rare or low-confidence features. Quality control summaries were generated in parallel. Filtered pathway tables were imported into R (v4.3.2) for TSS normalization on log transformed pathways and differential abundance analysis using Maaslin2 v1.10.0 (Mallick et al., 2021).

#### Behavioral correlations

Associations between behavioral sensitivity and microbial taxa were tested using MaAsLin2, modeling CLR-transformed abundances as a function of continuous sensitivity score with cohort and individual mouse id included as a random effect. P-values were corrected for multiple testing using the Benjamini–Hochberg procedure, and the significance cutoff was q < 0.05. Figures were generated using phyloseq (McMurdie and Holmes 2013), tidyverse (Wickham et al. 2019), and ggplot2 (Wickham 2011).

## Results

### One hour social interaction with a CFA-injected mouse induces the social transfer of pain in mice

Social interaction with a mouse experiencing hyperalgesia can induce matched pain behavior in bystander mice. (Smith et al. 2021) We replicated this in 2 cohorts of “pain” mice (n=8 per cohort, N=16) which were injected with complete Freud’s adjuvant (CFA) to induce inflammatory pain. (Figure 1A) Bystander animals were exposed to CFA-injected animals for one hour immediately after CFA injection. Compared to controls, CFA-injected animal mechanical sensitivity thresholds were significantly decreased at 1 (0hr) and 4 hours post-social interaction. (Kruskal-Wallis, Dunn post-hoc, p= 0.008). (Figure 1B) Bystander mechanical thresholds were also significantly lower than controls at both time points (Kruskal-Wallis, Dunn post-hoc, p= 0.035). At 24 hours post-injection, CFA-injected mice exhibited significantly reduced mechanical thresholds compared to controls (Kruskal–Wallis, Dunn post-hoc, p = 0.007), whereas thresholds in bystander mice were no longer significantly different from controls (p = 0.79) (Figure 1B). These findings indicate that bystanders initially develop hyperalgesia following social interaction with a mouse in pain, but this effect resolves in most bystanders by 24 hours.

**Figure 1:**
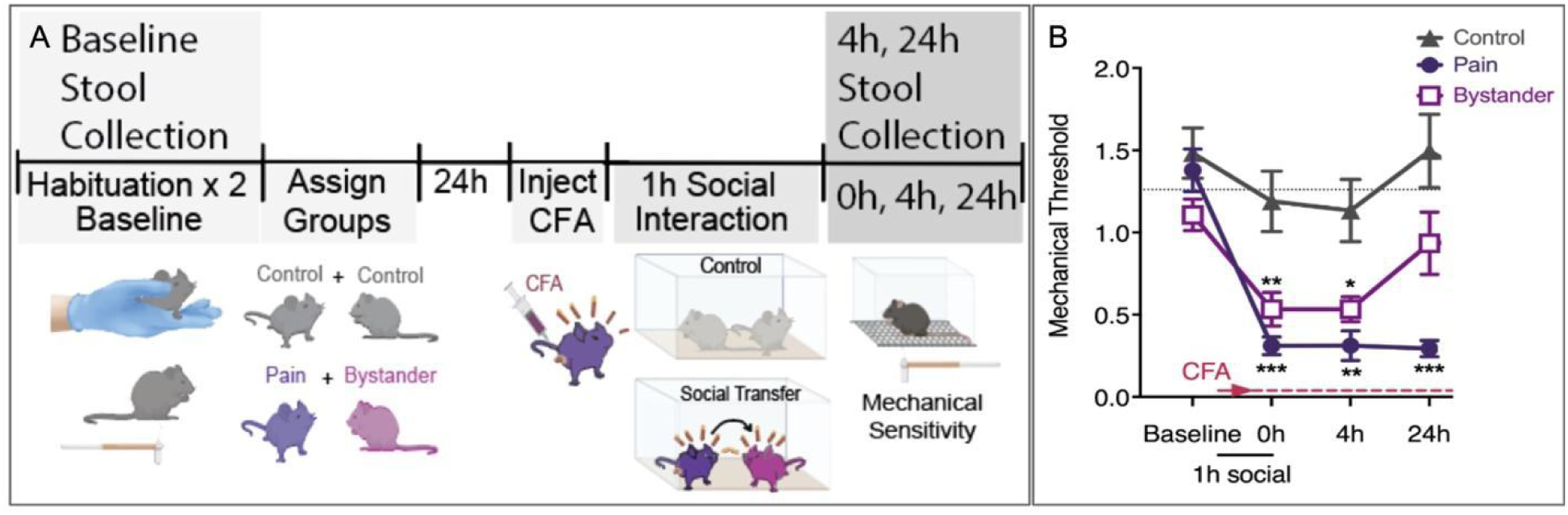
Social Transfer of Pain Confers Hyperalgesia to Bystander Mice. A: Experimental Setup; B: Mechanical Sensitivity testing using von Frey behavioral testing in two independent experiments of social interactions (n=24, N=48), comparing Von Frey behavioral testing for pain levels at 4 hours post-social interaction as well as at 24 hours post-social interaction across groups.

### CFA-induced pain and socially transferred pain are associated with fecal microbial community compositional changes

We hypothesized that the gut microbiome would be altered in CFA-injected pain mice, as well as altered in bystanders following the social transfer of pain. To test this, two independent cohorts of control, pain and bystander mice were run through the social transfer of pain (n=8 per group) and sequenced for analysis of the gut microbiome’s response using community composition metrics. (Figure 2) Changes in alpha diversity were calculated over time within each experimental group. (Shannon Diversity; Figure 2A) Linear mixed effect modeling revealed that Shannon Diversity significantly increased between baseline and 4 hours in both CFA-injected animals (t_stat_=4.08, p = 0.0003) and bystander animals (t_stat_=2.65, p = 0.013), but not in control animals (t_stat_=-0.42, p = 0.68). Shannon Diversity did not alter from baseline to 24 hours in CFA-injected animals (t_stat_=0.80, p = 0.43) and bystander animals (t_stat_=0.54, p = 0.59), but significantly decreased in control animals (t_stat_=-3.48, p = 0.002). We also compared Shannon Diversity across CFA-injected, control, and bystanders at each timepoint, but no difference was found between groups at any timepoint.

**Figure 2:**
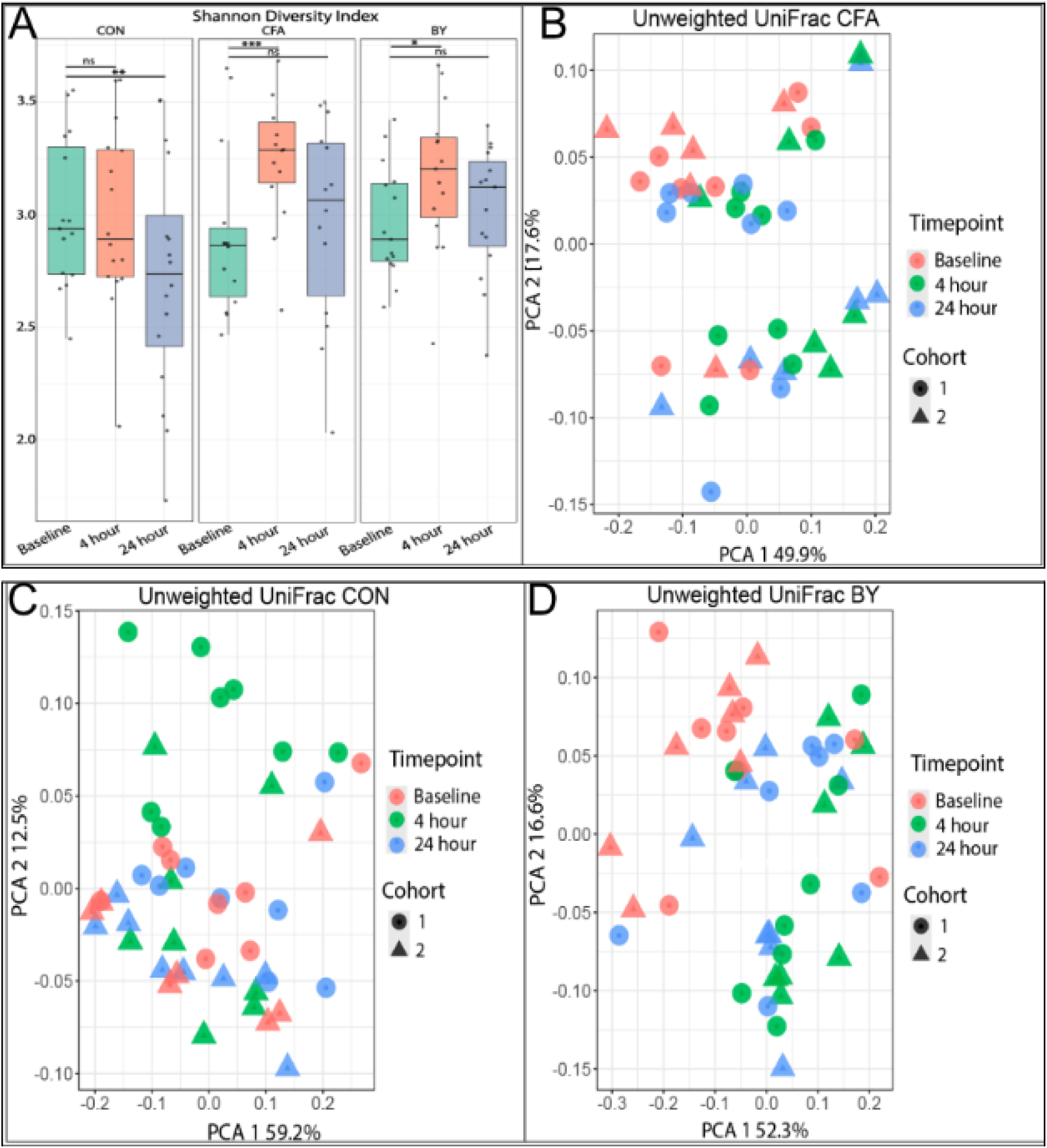
Pain and Social Transfer of Pain show Altered Community Composition from Controls. A. Shannon Diversity Index for Alpha Diversity measurements across sample groups and timepoints. (B) Unweighted Unifrac Principal Component Analysis for CFA-injected animals (CFA); (C), control (CON) animals; and (D) bystanders (BY).

We examined all the samples across time using Weighted and Unweighted Unifrac metrics and PERMANOVA analysis. (Supplemental Figure 1; Supplemental Table 2) Timepoint and behavioral cohort were found to be significant covariates for both metrics, while the experimental group was a significant covariate using Weighted UniFrac but not significant using Unweighted UniFrac. (p = 0.055) Weighted UniFrac PCA visualization suggested that samples cluster by cohort across time. (Supplemental Figure 1A) In Unweighted UniFrac PCA visualization, clusters were driven by presence or absence of unknown rare taxa that divided all samples into two distinct groups, regardless of experimental group or cohort. (Supplemental Figure 1B) The samples comprising the two distinct groupings alter over time, suggesting that animals gain (or lose) the taxa defining the groupings. No other covariates in this study could explain this grouping.

Comparing CFA-injected gut microbial samples to their own baseline, bacterial community composition differed significantly, as assessed by β-diversity, at 4 hours post-social interaction (5 hrs post injection; Pairwise PERMANOVA, Unweighted UniFrac, R² = 0.28, p = 0.004) as well as 24 hours post-social interaction (Pairwise PERMANOVA, Unweighted UniFrac, R² = 0.27, p = 0.005) (Figure 2B).

Community composition did not differ significantly between 4 hours post-social interaction and 24 hours post-social interaction in CFA-injected animals (Pairwise PERMANOVA, Unweighted UniFrac, R² = 0.22, p = 0.13) (Figure 2B). Similarly, bystander gut microbial community composition differed significantly from baseline at 4 hours post-social interaction (Pairwise PERMANOVA, Unweighted UniFrac, R² = 0.30, p = 0.001) and 24 hours post-social interaction (Pairwise PERMANOVA, Unweighted UniFrac, R² = 0.20, p = 0.033) (Figure 2C). However, no significant differences in community composition were observed between 4 hours and 24 hours post-social interaction (Pairwise PERMANOVA, Unweighted UniFrac, R² = 0.12, p = 0.26). In control samples, no differences in gut microbial community composition were observed between baseline and 4 hours post-social interaction (Pairwise PERMANOVA, Unweighted UniFrac, R² = 0.15, p = 0.14) or 24 hours post-social interaction (Pairwise PERMANOVA, Unweighted UniFrac, R² = 0.12, p = 0.29) (Figure 2D). While community composition appeared altered between 4 hours and 24 hours post-social interaction in control samples (Pairwise PERMANOVA, Unweighted UniFrac), repeated analysis separately by cohort revealed that neither cohort individually showed significant differences at any timepoint by either Weighted or Unweighted UniFrac, likely due to reduced power (Supplemental Table 1).

Taken together, these results indicate that gut microbial community structure and diversity are altered by CFA-induced pain, as well as by socially transferred pain in bystander mice.

### CFA-induced pain and socially transferred pain associate with overlapping but unique microbial community profiles

As alpha and beta diversity of the fecal microbiome were altered in CFA-injected animals as well as in bystanders, we were interested in identifying specific microbial taxa that contributed to these differences. Differential abundance analysis was performed using Maaslin2 on gut microbiome samples by comparing baseline samples to 4 hours post-social interaction as well as 24 hours post-social interaction at the species level, adjusting for behavioral cohort and repeated measures. (Figure 3) No taxa were identified as significantly altered in control animals at 4 hours or 24 hours post-social interaction when compared to baseline. At 4 hours post-social interaction, 13 taxa were found to be significantly altered compared to baseline in both CFA-injected and bystander animals. (Figure 3A) *Flavonifractor plautii* (*Oscillospiraceae)*, *Fusicatenibacter saccharivorans (Lachnospiraceae)*, and *Traorella massiliensis (Erysipelotrichaceae)* were all significantly increased in proportion, with effect sizes between 7 and 12. (Figure 3A) Ten species across families *CAG 272 (*Order*: Clostridiales)*, *Eggerthellaceae*, and *Muribaculaceae* were significantly decreased in proportion at 4 hours post-social interaction compared to baseline. (Figure 3A) However, by 24 hours post-social interaction, only 3 taxa which were altered in CFA-injected animals, and 4 taxa for bystander animals. (Figure 3A) Both CFA and BY microbiomes remained significantly depleted in Avispirillium sp 011957885 (Order: *Clostridiales*) and D16-63 sp003612475 (*Eggerthellaceae)*. (Figure 3A) However, only CFA-injected animals maintained an increased proportion of *F. plautii,* while bystander animals maintained an increased proportion of both *F. saccharivorans* and *T. massiliensis*.

**Figure 3:**
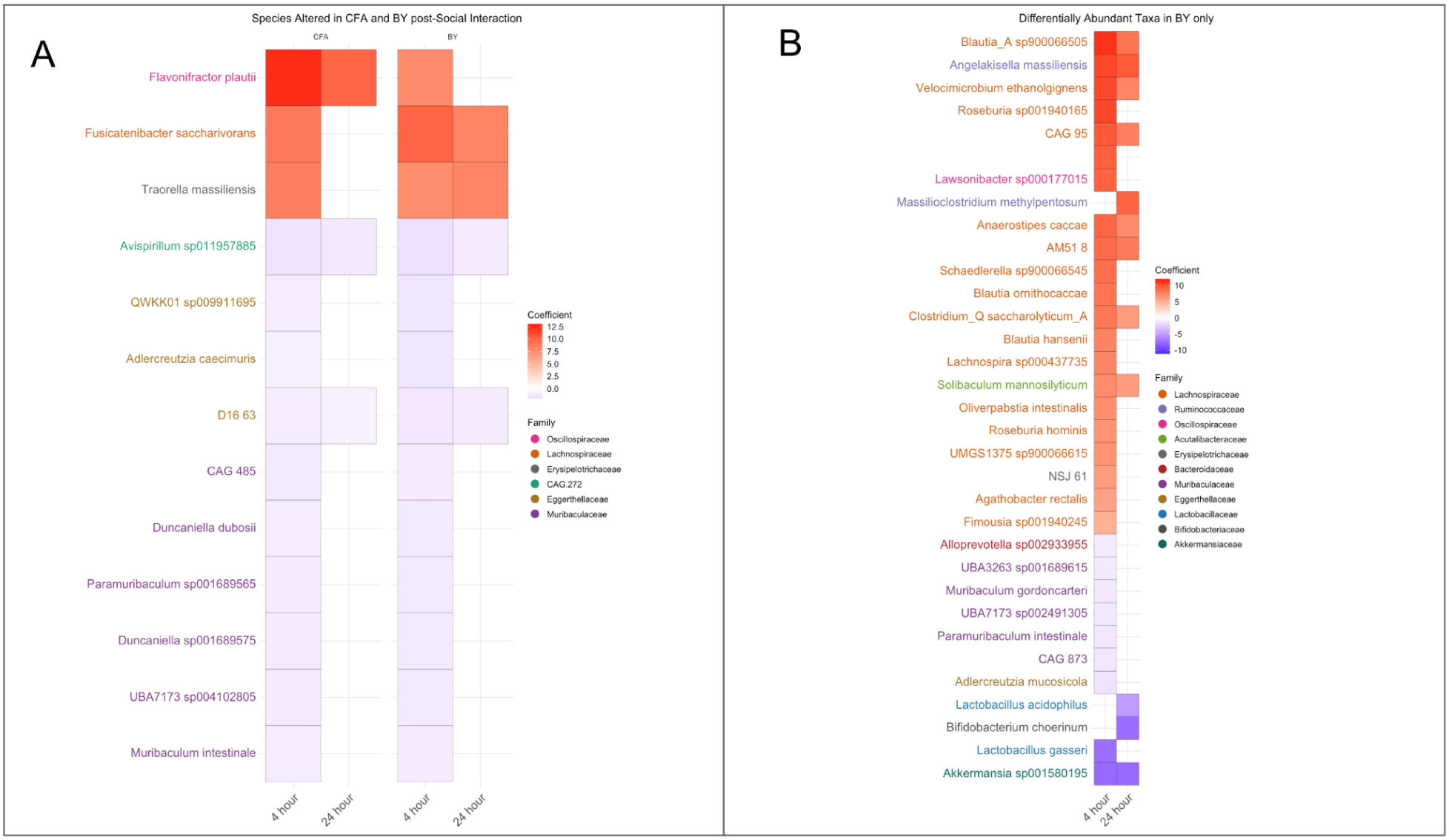
Differential Abundance Analysis of Gut Microbes in Pain and Social Transfer of Pain across Time. A) A heat map of all taxa identified as differentially abundant from baseline to 4 hours post-social interaction and 24 hours post-social interaction in species altered in both CFA-injected and bystander animals. Maaslin2 was used on CLR transformed data and a cutoff of q < 0.05. B) A heat map of species identified as differentially abundant from baseline to 4 hours post-social interaction and 24 hours post-social interaction that were uniquely altered in bystander. Maaslin2 was used on CLR transformed data and a cutoff of q < 0.05.

The bystander gut microbiome showed changes in the proportion of over 30 additional species at 4 hours post-social interaction which were not altered in CFA-injected animals. (Figure 3B) Of note, 13 of these species were within the family *Lachnospiracae*, a family which was also increased in CFA-injected animals. Species in genera *Blautia*, *Roseburia*, and *Clostridium* were all significantly increased in proportion with effect sizes from 5-11. (Figure 3B) Meanwhile, 9 bacterial species were significantly decreased in proportion, including 5 species from *Muribaculaceae,* a family also significantly decreased in CFA animals. *Lactobacillaceae gasseri*, *Akkermansiaceae sp 001580195,* and *Adlercruzia mucosicola* were also significantly decreased in proportion at 4 hours post-social interaction. (Figure 3B) At 24 hours post-social interaction, 12 species were significantly altered compared to baseline in bystanders. Nine overlapped with the 4 hours post-social interaction profile which were annotated to families *Lachnospiraceae*, *Ruminococcaceae*, and *Akkermansiaceae*. Three were uniquely altered at 24 hours post-social interaction in bystander animals, *Massilioclostridum methypentosum (Ruminococcaceae), Lactobacillus acidophilus*, and *Bifidobacteriaceae choerinum.* (Figure 3B)

Using Maaslin2 linear mixed effect modeling adjusted for cohort and repeated measures, we also explored associations between individual species of bacteria and behavioral measurements across all samples. (Supplemental Figure 3: Supplemental Table 3) At q < 0.05, *Clostridium Q.saccharolyticum_A* (Family: *Lachnospiraceae*) negatively correlated with sensitivity scores (coef =-2.13, q = 0.01), and *Akkermansia sp001580195* positively correlated with sensitivity scores (coef=2.35, q = 0.01). (Supplemental Table 3) Both of these taxa were significantly altered in bystanders at 4 hours and 24 hours. (Figure 3B) Interestingly, these taxa were not significantly altered in CFA animals. (Figure 3A)

Taken together, these results suggest the gut microbiome of bystanders responds to the social transfer of pain, and that the response partially mirrors the changes seen in the gut of animals with CFA-induced pain. The unique response of several microbes particularly in the *Lachnospiraceae* family within bystanders may indicate differentiation between animals experiencing CFA-induced pain and those receiving socially transferred pain.

### Predicted microbial functional changes are similar post CFA-induced pain and social transfer of pain

To gain insights on the metabolic potential of the gut microbial response to pain and social transfer of pain, we functionally annotated metagenomic reads to assess the proportions of predicted functional pathways and used Maaslin2 to perform differential abundance analysis within CFA-injected, bystander, and control groups. (Figure 4) No pathways were identified as significantly altered in control animals at 4 hours or 24 hours post-social interaction when compared to baseline. (Supplemental Table 4) When compared to baseline, both CFA-injected and bystander animals demonstrated altered functional potential for 24 pathways at 4 hours and 24 hours post-social interaction. (Figure 4A) Overall, pathways showing significantly increased proportions in both CFA-injected and bystander groups were related to alternative carbon/nitrogen sourcing (degradation of ascorbate, sugars, and purine) and one-carbon metabolism (folate transformation/polyglutamylation). Several pathways related to acetate and butyrate production were also found to be enriched across CFA-injected and bystander animals, including pyruvate fermentation to butanoate and L-lysine fermentation to acetate and butanoate. Both CFA-injected and bystander animals also demonstrated significantly decreased functional potential for carbohydrate metabolism (sucrose degradation, heterolactic fermentation), as well as amino acid biosynthesis.

**Figure 4:**
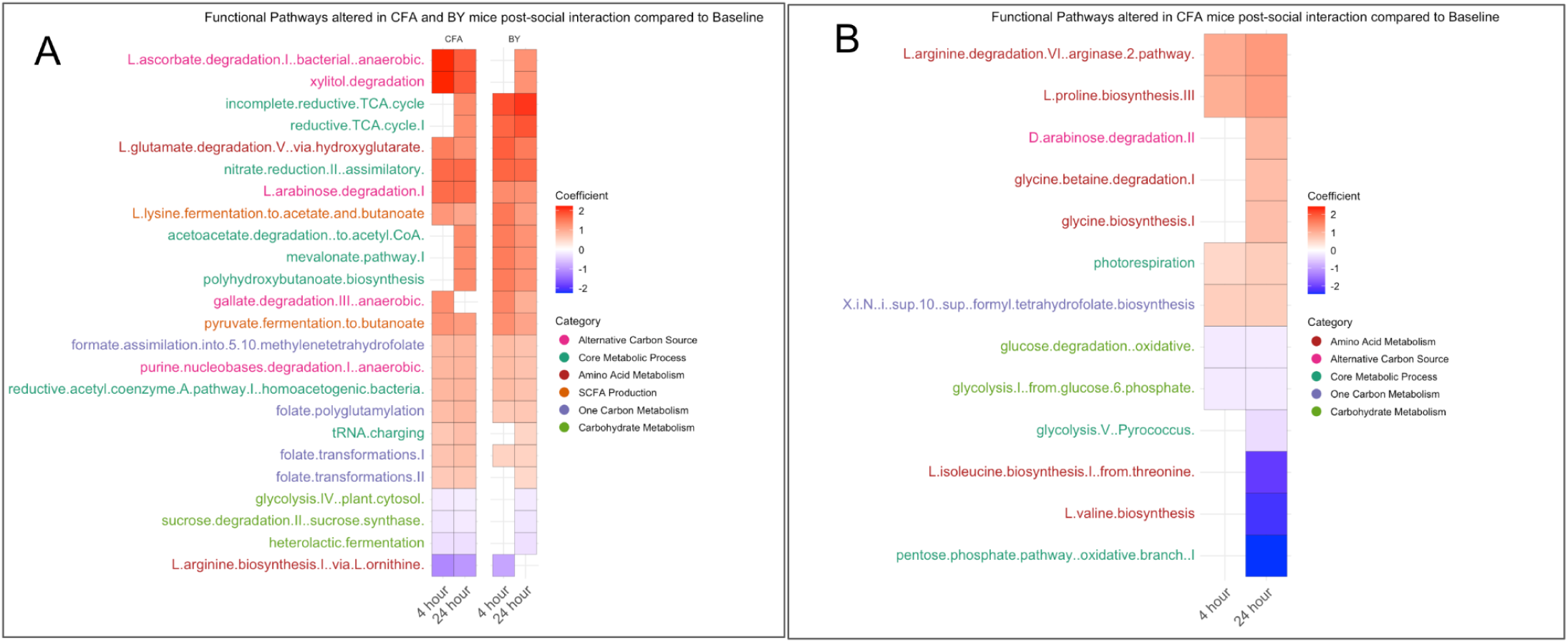
Functional Potential of the Gut Microbiome was Altered after Pain and Social Transfer of Pain. Metagenomic reads were functionally annotated and timepoints 4 hours and 24 hours post social interaction were compared to baseline with CFA-injected animals and bystander animals statistically to determine alterations in functional potential of the fecal microbiome using Maaslin2 differential abundance at q < 0.05.A) Functional pathways overlapping in both CFA and BY animals. Text is colored by functional category groupings. B) Functional pathways identified as significantly altered exclusively in CFA-injected animals.

At 4 and 24 hours post-social interaction, CFA-injected gut microbiomes showed changes in proportion of an additional 13 functional pathways when compared to baseline that were not found to be altered in bystander animals. (Figure 4B) Unique microbial metabolic pathways in CFA-injected animals related to carbohydrate metabolism (glucose degradation, glycolysis) were significantly reduced in proportion.

Increased proportions for proline and glycine biosynthesis and decreases for isoleucine and valine biosynthesis were also observed. (Figure 4B) Of the 13 altered functional pathways, half were found to be altered only at 4 hours post-social interaction, while all 13 were found to be altered at 24 hours post-social interaction. Bystander animal gut microbiomes only had 6 uniquely altered pathways. (Supplemental Figure 2) At 4 hours but not 24 hours post-social interaction, bystander animals demonstrated a significant decrease in the functional potential for the degradation of super oxide radicals. Key metabolic processes like the Calvin-Benson Bassham cycle and alternative sugar degradation (allose) showed significantly increased proportions. Overall, these results indicate that the functional potential of the microbiome is altered similarly by both CFA-induced pain and socially transferred pain, with increases in microbial potential to utilize alternative carbon sources and produce short chain fatty acids, and decreased potential for carbohydrate metabolism and amino acid biosynthesis.

## Discussion

This study investigated the effects of the social transfer of pain on the gut microbiome of mice. Though pain is a well-characterized modulator of the gut microbiome, to date no research has focused on elucidating the relationship between socially induced pain experiences and the gut microbiome. This study represents a preliminary characterization of gut microbiome alterations post-social transfer of pain. Specifically, we wanted to understand if the bystander gut microbiome was altered during the experience of hypersensitivity. We were also interested in understanding if any gut microbial community compositional changes in bystanders would resemble those observed in the CFA-injected animals. To this end, we collected fecal samples from CFA-injected animals and bystanders animals prior to as well as 4 hours and 24 hours post-social interaction and measured taxonomic changes in alpha diversity, beta diversity as well as differential abundance of taxonomic and functional potential features.

Bystander animals were exposed to animals injected with CFA for one hour, which successfully induced the social transfer of pain in bystanders at 4 hours post social interaction represented by mechanical hypersensitivity. Stool samples collected from bystander animals at this timepoint revealed the gut microbiome community composition was significantly altered when compared to baseline. Differential abundance analysis indicated 43 species showed a significant shift in proportion. Bystanders uniquely exhibited a bloom in the proportion of 17 species across 12 genera within *Lachnospiraceae*, as well as decreases of *Lactobacillus gasseri* and *Akkermansia sp 001580195* at 4 hours post-social interaction. The functional potential of the gut microbiome was also significantly altered towards utilization of alternative sources for carbon and fermentation production of butyrate and acetate. Though no studies characterize the impact of socially induced pain on the gut environment, these preliminary insights align with the taxonomic and functional gut microbial profile of a response to pain. (Ma et al. 2020; Gonzalez et al. 2025) When a host experiences noxious stimuli, a systems-level response occurs that may alter the gut environment through changes in nutrient availability, oxygenation, and immune signaling, thus creating redox and nutrient conditions that favor fermentative guilds, nitrate respiration, and folate-dependent one-carbon flux, while disfavoring saccharolytic commensals and anabolic metabolism. (Ustianowska et al. 2022; Agirman et al. 2021; Zeng et al. 2017; Winter et al. 2013; Engevik et al. 2019; Qiao et al. 2020) Our results thus suggest that socially transferred pain may alter the gut environment towards conditions that support a stress-tolerant gut microbial community.

We were also interested in understanding if the gut microbial response of bystanders resembles the CFA-injected mice microbiome post-social interaction. At the 4-hour timepoint, all 13 microbial species that showed altered proportions in CFA-injected animals were also altered in the same direction and at similar magnitude in the bystanders. Further, CFA-injected and bystander animals shared 24 functional metabolic pathways that altered post-social interaction. This similar microbial response suggests that the physiological features that shape the gut microbiome of CFA animals may be similar in bystanders.

Importantly, outside of the overlap, bystander animals and CFA-injected animals had some unique taxonomic and predicted functional differences. The bystander bloom of Lachnospiraceae spp. at 4 h post-social interaction, together with decreased predicted capacity for reactive-oxygen-species handling (e.g., superoxide detox), is consistent with subtle mechanistic differences in host physiology during socially transferred pain, potentially involving shifts in epithelial oxygenation/redox state (Rivera-Chávez et al. 2017; Imlay 2013; Seixas et al. 2021) and the availability of fermentable carbon substrates upon which Lachnospiraceae rely (Vacca et al. 2020; Zaplana et al. 2023).

Our research provides preliminary insights on the gut microbial response effects of socially transferred pain induced through social interaction between a bystander and a mouse injected with CFA. However, we only explored one mechanism of inducing pain in the “donor” animals. Socially transferred pain may also be induced via alcohol or morphine withdrawal, capsaicin, or formalin (Smith et al. 2016, 2021), so it will be important to understand if these other mechanisms lead to similar differences. Moreover, experiences like analgesia, fear, and inflammation can be socially transferred that may also uniquely alter the gut microbiome. Future studies should address this research gap by exploring the response, if any, to the social transfer of these experiences. As the mechanism for social transfer remains unclear, we explored the possibility that coprophagy between bystanders and CFA-injected animals during the social interaction may lead to similar changes in the gut microbiome. Visual footage of the one-hour social interaction did not reveal any instances of coprophagy, but future studies should explore this variable.

Finally, we only tested our results in male mice. Future studies could examine if the social transfer of pain differentially alters the guts of female mice.

## Ethics

All procedures were approved by the Institutional Animal Care and Use Committee at University of California San Diego (Protocol No. S22191) and were conducted in accordance with the National Institutes for Health Guidelines for the Care and Use of Laboratory Animals and the Guidelines for the Care and Use of Mammals in Neuroscience and Behavioral Research.

## Data availability

Data will be made available upon peer-reviewed publication.

## Acknowledgements

This publication includes data generated at the UC San Diego IGM Genomics Center utilizing an Illumina NovaSeq 6000 that was purchased with funding from a National Institutes of Health SIG grant (#S10 OD026929).

## Supplemental Materials

Supplemental tables are provided as a single spreadsheet file, with each table presented as a separate tab, in a separate file. Supplemental Figures are provided in a separate file.

### Supplemental Table Captions

Supplemental Table 1. Cohort-level analysis of beta diversity for control animals. Animals were subsetted by cohort 1 or cohort 2, and pairwise PERMANOVA analyses were performed per cohort. R2 values and p values are reported.

Supplemental Table 2. Weighted and Unweighted Unifrac PERMANOVA analyses were performed globally across samples. R2 values and p values are reported.

Supplemental Table 3. Correlation between mechanical sensitivity scores and clr-transformed abundances of microbial taxa at the species level. Maaslin2 was used for linear mixed effect modeling adjusted for cohort and repeated measures. Q values < 0.05 were used in the main text.

Supplemental Table 4. Differential abundance of predicted functional pathways in control animals. Metagenomic reads were functionally annotated and Maaslin2 was used to determine enrichment or depletion in relative abundance of pathways. Control animal values are shown per pathway.

