## Supplemental Figures for "My Gut Feels Your Pain - The Social Transfer of Pain Remodels the Gut Microbiome"

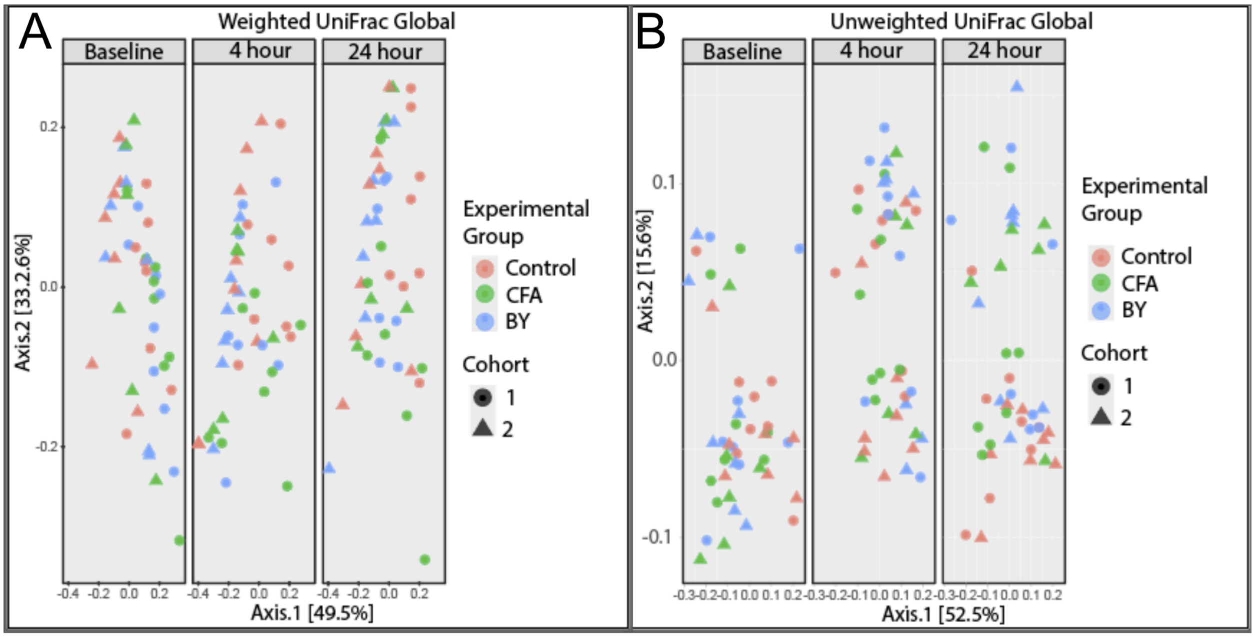


Supplemental Figure 1, Beta Diversity of All Samples Across Time. Weighted (A) and Unweighted (B) UniFrac Principal Component Analysis. Graphs are colored by experimental group and behavioral cohort is represented by shape.

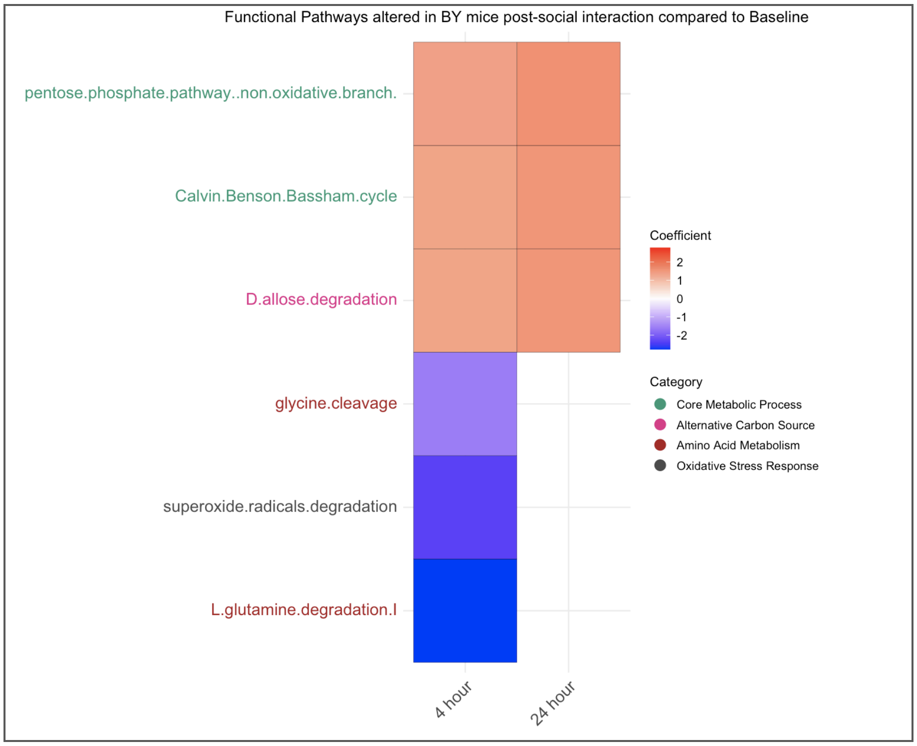


Supplemental Figure 2, Functional Potential of the Gut Microbiome is Altered Post-Social Interaction in Bystanders. Metagenomic reads were functionally annotated and timepoints 4 hours and 24 hours post-social interaction were compared to baseline within the bystander group to statistically determine alterations in functional potential of the fecal microbiome using Maaslin2 differential abundance at $q < 0.05$. Text is colored by functional category groupings. Pathways uniquely altered in bystanders are shown.

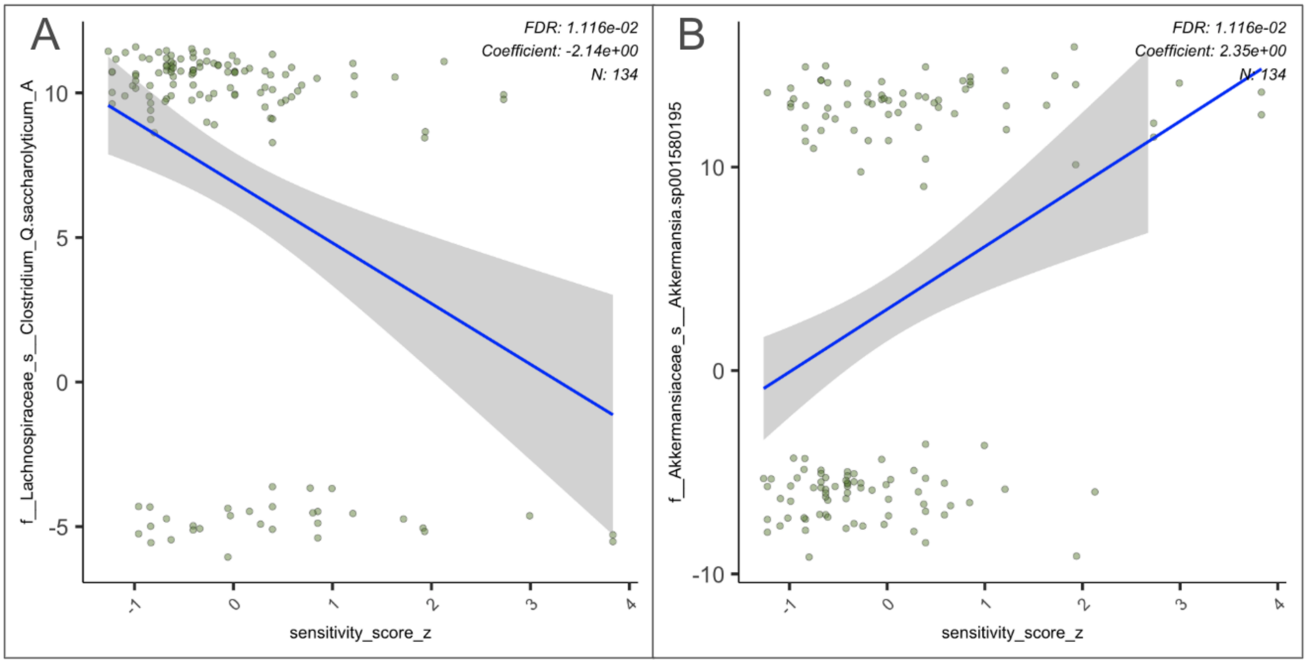


Supplemental Figure 3, Correlation between mechanical sensitivity score and abundance. Abundance data was center log transformed and correlated to mechanical sensitivity scores across all experimental group and timepoints. Mechanical sensitivity score is inversely related to pain experienced. A) Correlation of mechanical sensitivity scores with *Clostridium Q saccharolyricum A* (Family: *Lachospiracaeae*). B) Correlation of mechanical sensitivity scores with *Akkermansia sp 001580195* (Family: *Akkermansiaceae*)
